# Prolonged visual deprivation in adults induces non-homeostatic plasticity of feed-forward excitation from visual thalamus to cortex

**DOI:** 10.1101/2024.12.04.626829

**Authors:** Sachiko Murase, Daniel Severin, Louis Dye, Lukas Mesik, Cristian Moreno, Alfredo Kirkwood, Elizabeth M. Quinlan

## Abstract

Age constrains plasticity at inputs from first order thalamic nuclei to the cortex, endowing stability to ascending, feed-forward projections. However, here we show that prolonged visual deprivation can induce robust and reversible plasticity at thalamocortical synapses in layer 4 pyramidal neurons in the adult mouse primary visual cortex. The plasticity engaged by prolonged visual deprivation is non-homeostatic and mediated by changes in presynaptic function.

## Introduction

Synaptic plasticity is constrained by age, endowing mature neuronal circuits with enhanced stability. However, robust receptive field plasticity can be rejuvenated in the adult rodent visual cortex by visual deprivation through prolonged dark exposure (DE) and subsequent light reintroduction (LRx) (He et al., 2006, 2007). In amblyopic rats, visual stimulation following dark exposure drives the recovery of visual acuity through the amblyopic eye (He et al., 2007; Montey and Quinlan, 2011; Eaton et al., 2016). The rejuvenation of plasticity by DE has been also demonstrated in felines (Duffy et al., 2016) and mice (Stodieck et al., 2014; Murase et al., 2017; Jeon et al., 2022). However, our understanding of the mechanisms that couple prolonged sensory deprivation with the rejuvenation of plasticity is incomplete.

Alterations in synaptic efficacy can serve to amplify or compensate for modifications in synaptic inputs (non-homeostatic/homeostatic, respectively). Binocular visual deprivation induces well-documented homeostatic changes at cortico-cortical synapses in the primary visual cortex (Wen and Turrigiano, 2024). DE increases the expression of the GluN2b subunit of the NMDAR in adult rodent V1 (He et al., 2006), endowing NMDARs with high agonist affinity, high open probability and slow deactivation kinetics. High GluN2b levels are associated with a low induction threshold for correlation-based plasticity (Cooper and Bear, 2012; Lee and Kirkwood, 2019). Accordingly, the increase in EPSC amplitudes observed in V1 pyramidal neurons following DE is blocked by antagonism of the GluN2b receptor (Bridi et al., 2018). The homeostatic increase in mEPSC amplitudes in superficial cortical lamina following visual deprivation is expressed in juveniles and adults (Goel et al., 2006; Goel and Lee, 2007). In humans, brief DE (60 min) is sufficient to increase the local concentration of glutamine/glutamate in human adult V1, suggesting a rapid homeostatic response (Min et al., 2023).

In contrast, synapses between first order sensory thalamus and cortex lose plasticity early in postnatal development. Homeostatic regulation of mEPSC amplitudes following DE is not observed at thalamocortical synapses (Petrus et al., 2014, 2015; Chokshi et al., 2019). Classic experiments in the rat barrel cortex demonstrate that pairing-induced LTP, NMDAR-dependent LTD and silent synapse conversion are lost early at TC synapses (Crair and Malenka, 1995; Isaac et al., 1997; Feldman et al., 1998). Similarly, in the primary visual cortex, LTP in layer 4 neurons induced by stimulation of white matter is constrained prior to plasticity at L4-L2/3 synapses (Jiang et al., 2007). In cat V1, mesoscopic anatomical plasticity of geniculocortical afferents induced by monocular deprivation (MD) is lost early in development while robust plasticity persists at cortico-cortical synapses ((Wiesel and Hubel, 1963; Antonini and Stryker, 1993, 1996). This work suggests that homeostatic and non-homeostatic mechanisms are lost with age at thalamocortical (TC) synapses.

However, plasticity at thalamocortical synapses can be induced at spared inputs following extraordinary manipulations, such as deafferentation. For example, transection of the branch of the trigeminal nerve that innervates the whisker pads results in potentiation of thalamic inputs to the spared barrel cortex (Yu et al., 2012; Chung et al., 2017; Jie et al., 2025). Similarly, visual deprivation induces cross-modal strengthening of thalamocortical (TC) synapses in auditory cortex, but does not impact TC synapses in primary visual cortex (V1; Petrus et al., 2014). Environmental enrichment, which can promote plasticity throughout the lifespan (Sale et al., 2007) may induce sprouting of dLGN axons (Mainardi et al., 2010).

Contrary to expectations, our previous work identified thalamocortical synapses as a potential locus for the rejuvenation of plasticity in the primary visual cortex by dark exposure and light reintroduction (Montey and Quinlan, 2011; Murase et al., 2017). Here we used a series of orthogonal experimental approaches to specifically ascertain if DE/LRx can modify the efficacy of feed-forward synapses from first order thalamus to visual cortex in adult mice. We find that prolonged DE induces significant changes in presynaptic structure and function of TC synapses, including changes in ultrastructure, molecular geometry, visually-evoked calcium magnitudes and neurotransmitter release probability. The presynaptic plasticity engaged by prolonged visual deprivation at TC synapse was reversible and non-homeostatic, thereby serving to relay and amplify changes in feed-forward afferent drive to the visual cortex.

## Results

To ask if prolonged DE regulated the efficacy of feed-forward synapses from first order thalamus to L4 in the adult mouse visual cortex, we first optogenetically isolated the activity of thalamic axons in acute slices of primary visual cortex (Fig. 1a, b). AAV5-CaMKIIa-hChR2(H134R)-mCherry was delivered to dLGN of adult C57BL/6J mice (>postnatal day (P) 90). Optogenetically evoked EPSCs at the light intensity that evoked a response in 50% of trials (470 nm, 3 msec) were measured in V1b L4 neurons in TTX and 4-AP following desynchronization of vesicle release by substituting Sr^2+^ for Ca^2+^ (Dodge Jr. et al., 1969; Petreanu et al., 2009; Whitt et al., 2022). In control normal-reared adult mice (NR, raised in a 12/12 h L/D cycle), light-evoked EPSCs had the expected mean amplitude of ∼13 pA, which was unchanged following 10 days of DE (Fig. 1c, d). Brief LRx (2 h) induced a small, but significant decrease in Sr^2+^-mEPSC amplitudes (mean±sem: 12.9±0.7 pA; DE 13.0±0.6 pA; LRx 11.2±0.4 pA; n = 31, 26, 25 cells from 6, 5, 5 subjects, respectively, Kruskal-Wallis H (2) = 10.39, p = 0.0055, *post hoc* Dunn’s multiple comparison test; Fig. 1c, d). Thus, the estimated quantal amplitude of visually-evoked TC activity is relatively stable.

**Fig. 1.**
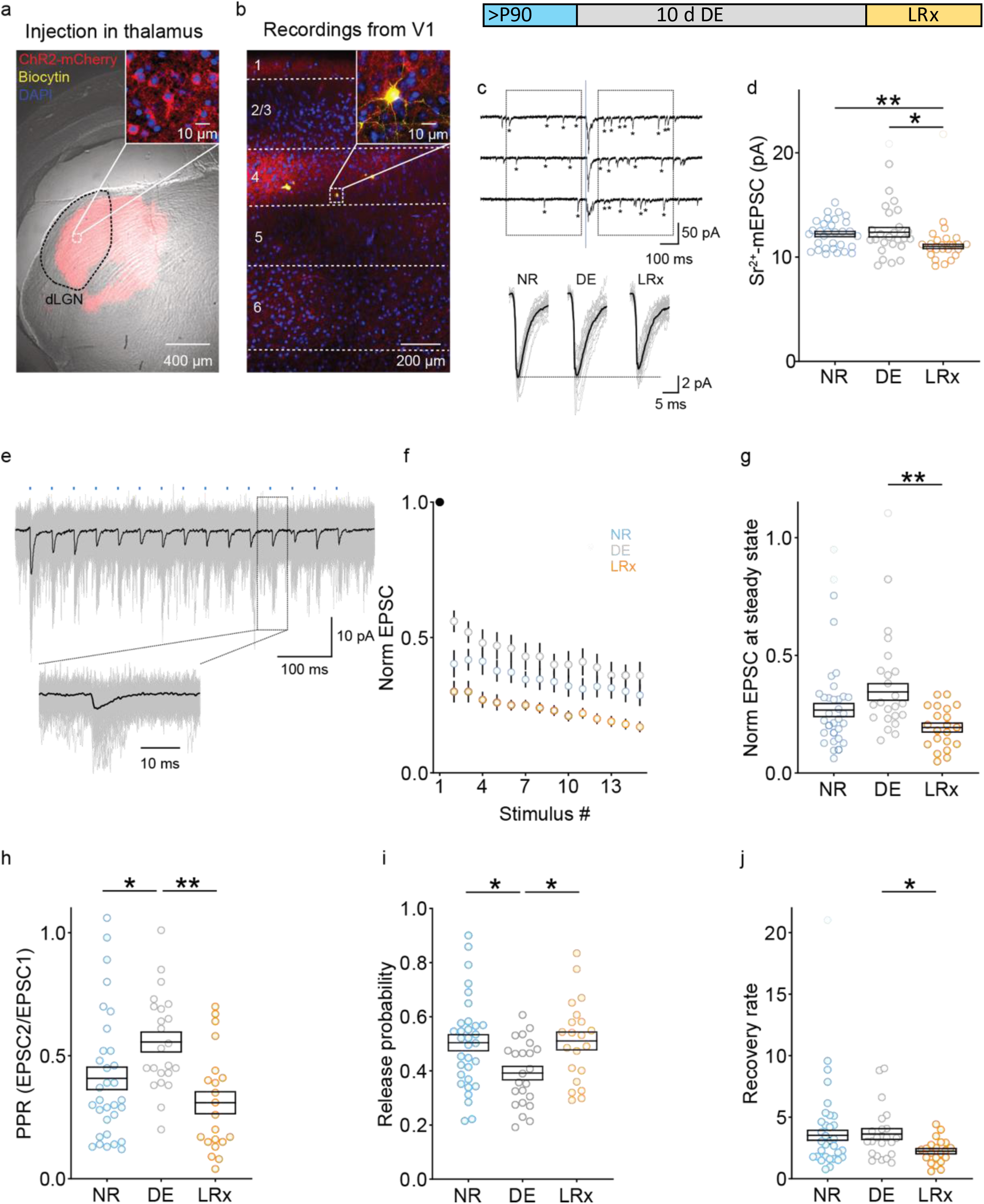
DE/LRx inducse bidirectional changes in presynaptic function of thalamocortical synapses in adult mouse V1b L4. a) AAV-CaMKIIa-hChR2-mCherry delivered to the dLGN (red) effectively transfected thalamic axons. An overlay of the bright field and confocal images reveals the viral injection site in thalamus (mCherry). The inset shows dLGN neurons counterstained with DAPI (blue) and expressing ChR2-mCherry. b) Example of a pyramidal neuron in layer 4 V1b filled with biocytin during whole cell recording (yellow). Note mCherry-positive axonal processes are enriched in L4. c) Inset: Experimental timeline. Adult C57B/L6 mice (>P90) received 10 days of DE or DE followed by 2 h of LRx. Top: Representative examples of light-evoked Sr^2+^-desynchronized miniature EPSCs (Sr^2+^-mEPSCs, 470 nm, 3 msec) in layer 4 V1b. Boxes = 400 ms time windows used for collecting Sr^2+^-mEPSCs (asterisks) before and after LED stimulation to evoke release (blue line; 3 ms duration). Bottom: Average traces for all the cells shown in Fig. 1d (gray) and average traces for all neurons in all subjects (black) for NR, DE, and LRx. d) Sr^2+^-mEPSCs were unchanged by DE (grey) and decreased after LRx (2 hr, yellow; mean Sr^2+^-mEPSC amplitude ± sem: NR (blue), 12.9±0.7 pA; DE 13.0±0.6 pA; LRx 11.2±0.4 pA; n = 31, 26, 25 cells from 6, 5, 5 subjects, respectively, Kruskal-Wallis H (2) = 10.39, p < 0.01, *post hoc* Dunn’s multiple comparison test). e) Repetitive optogenetic stimulation in the absence of Sr^2+^ (15 pulses of 470 nm LED for 3 msec at 25 Hz (blue dashes) at intensity that evoked a response in 100% of trials). Inset – higher magnification of boxed recording snippet. f) EPSCs normalized to the response to the first stimulus (filled dot). Visual experience modifies the response to repetitive stimulation (n = 32, 23, 21 cells from 8, 6, 5 subjects for NR, DE, LRx, respectively. Repeated measures ANOVA, F = 63.3, **p < 0.01, *post hoc* Bonferroni). g) Normalized EPSC at steady state. h) Paired pulse ratio (EPSC in response to stimulus 2/stimulus 1) is increased by DE and recovered by LRx. i) Presynaptic release probability is decreased by DE and recovered by LRx. j) LRx decreases synaptic vesicle recovery rate. *p < 0.05, **p < 0.01 for *post hoc* tests.

In contrast, DE/LRx induced significant changes in neurotransmitter release probability in response to repetitive stimulation of TC synapses. Optogenetically-evoked AMPAR-dependent EPSCs were induced in L4 neurons by LED trains (Fig. 1e, 15 repetitions at 25 Hz at the light intensity that evoked a response in 100% of trials in 100 μM APV and 20 μM bicuculine). Normalized TC EPSC amplitudes reveal a significant depression of TC transmission in response to repetitive stimulation in all experimental conditions (Fig. 1f). However, the activity-dependent depression of TC synapses was significantly reduced by DE and increased LRx (Two-way ANOVA; visual experience F (2, 1095) = 71.36, p<0.0001, *post hoc* Bonferroni; interaction F (28, 1095) = 0.5042, p=0.9857; n = 32, 23, 21 cells from 8, 6, 5 subjects, respectively; Fig. 1f). Normalized approximate steady state amplitude, calculated as the mean amplitude of 11^th^ to 15^th^ EPSCs, was comparable to NR following DE and LRx (NR = 0.27±0.03, DE = 0.34±0.04, LRx = 0.19±0.02; n = 32, 23, 21 cells from 8, 6, 5 subjects respectively; Kruskal-Wallis H (2) = 11.22, p<001; *post hoc* Dunn’s multiple comparison test; Fig. 1g). The paired pulse ratio (PPR) calculated from the response to the second relative to the first stimulus pulse significantly increased following DE and decreased following LRx (NR = 0.41±0.05, DE = 0.56±0.04, LRx = 0.31±0.04; n = 32, 23, 21 cells from 8, 6, 5 subjects respectively; Kruskal-Wallis H (2) = 13.97, p = 0.0009, *post hoc* Dunn’s multiple comparison test; Fig. 1h). Accordingly, the presynaptic vesicle release probability was significantly decreased by DE and increased by LRx (NR = 0.50±0.03, DE = 0.39±0.02, LRx = 0.51±0.03; n = 32, 23, 21 cells from 8, 6, 5 subjects, respectively; One-way ANOVA, F = 4.699, p = 0.012, post hoc Tukey; Fig. 1i; as in Bridi et al. 2020 based on Wesseling and Lo, 2002). There were no significant changes in the vesicle recovery rate by DE or LRx (NR = 3.5±0.40, DE = 3.6±0.44, LRx = 2.2±0.22; n = 32, 23, 21 cells from 8, 6, 5 subjects respectively, Kruskal-Wallis H (2) = 6.838, p = 0.0327, *post hoc* Dunn’s multiple comparison test; Fig. 1j). Thus, prolonged visual deprivation and light reintroduction induce an unexpected, reversible regulation of presynaptic function at adult TC synapses.

The molecular and structural organization of axonal boutons are key determinants of presynaptic function. We therefore quantified fundamental parameters of the ultrastructure of thalamocortical synapses in transmission electron micrographs following DE and LRx. Images were collected from V1 layer 4 (at 360±40 μm from dura). All synaptic profiles with high contrast boundaries delineating the presynaptic membrane, active zone, post-synaptic membrane and post synaptic density were quantified.

Presynaptic area is an established predictor of presynaptic origin (Nahmani and Erisir, 2005). Accordingly, the distribution of presynaptic area was bimodal across experimental conditions (n = 264, 219, 320 synapses for Con, DE, LRx, respectively; Fig. 2a). Synapses in the top 15^th^ percentile of presynaptic area had larger cleft widths and PSD lengths, as predicted for TC synapses (cleft width: 21.6±0.3 nm vs. 19.8±0.1 nm; PSD length: 472±30 nm vs. 350±8 nm, p<0.01, Student’s T-test; n=230, 34). Larger synapses also had more synaptic vesicles in contact with the presynaptic membrane (“docked” synaptic vesicles; TC = 4.63±0.37, CC = 3.11±0.12, p<0.01, Student’s T-test) and a larger presynaptic area occupied by synaptic vesicles (synaptic vesicle zone, SVZ, TC = 0.26±0.2 μm^2^, CC = 0.16±0.01 μm^2^, p<0.01, Student’s T-test).

**Fig. 2.**
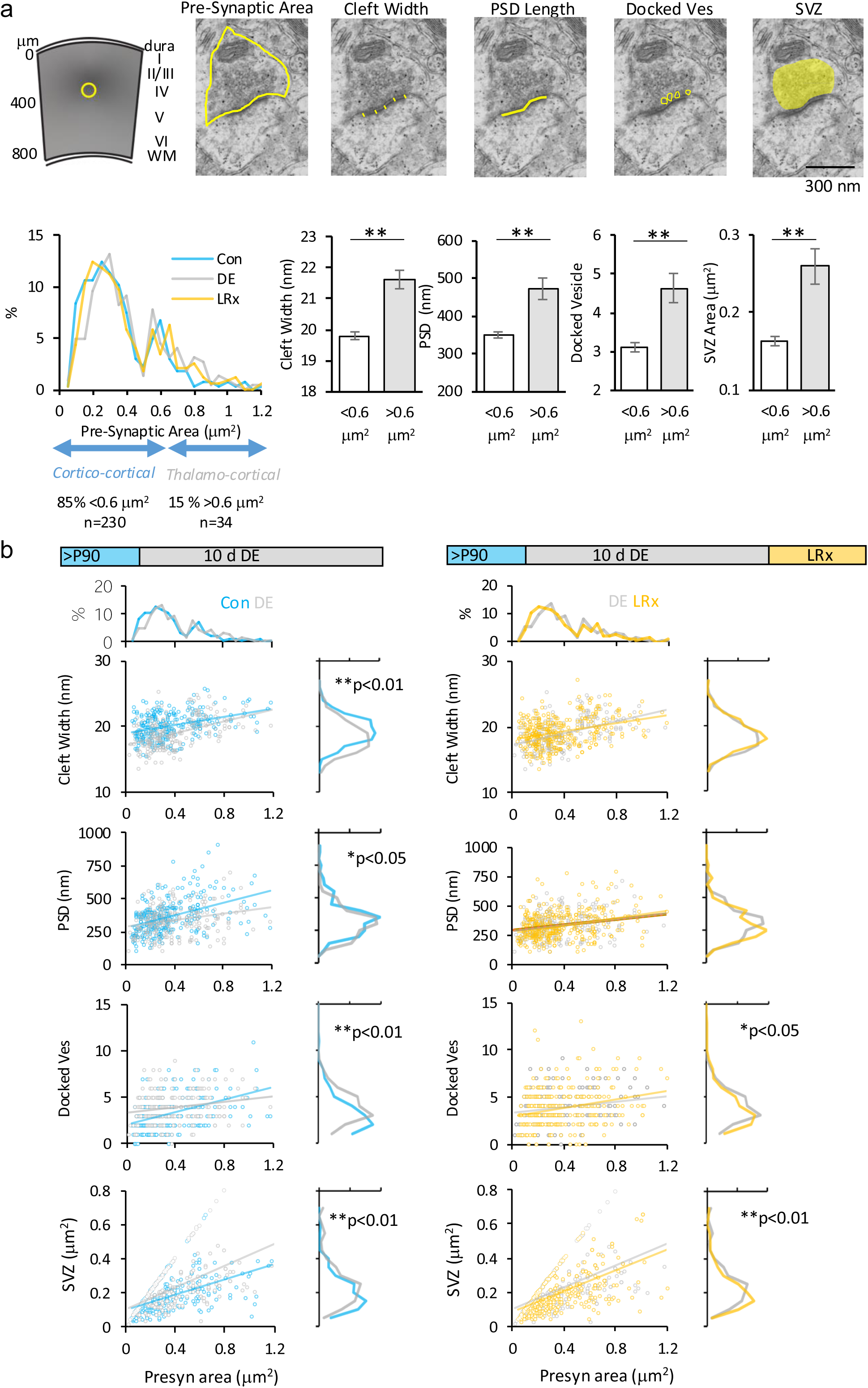
DE/LRx modify the ultrastructure of excitatory synapses in layer 4 of adult mouse V1b. a) Overlays depict sampling area in layer 4 and the structure quantified for presynaptic area, cleft width, PSD length, docked vesicles and synaptic vesicle zone. The distribution of synaptic profiles by presynaptic area reveals two peaks in all experimental groups (Con; blue, DE; grey, LRx; yellow): the larger population (15% of total; grey) has significantly larger cleft widths, PSD lengths, number of docked vesicles and synaptic vesicle zones (Student’s T-test, **p<0.01). b) Experimental timelines. Distribution of each ultrastructural parameter (Y axis) relative to presynaptic area (X axis). DE (grey) induces a significant difference in synaptic cleft width, PSD length, the number of docked vesicles and SVZ. LRx (yellow) induces a significant difference in the number of docked vesicles and SVZ (n = 264, 219, 320 synapses from 5, 4, 6 subjects for Con (blue), DE and LRx, Mann-Whitney Test, *p < 0.05, **p < 0.01).

A bivariate analysis was used to determine the impact of DE and LRx on each of these ultrastructural variables as a function of presynaptic area. Dark exposure induced widespread changes in ultrastructure, including a significant decrease in cleft width, notable in smaller synapses (Con = 20.0±0.1 nm, DE = 19.0±0.2 nm; p<0.01, Mann-Whitney Test, Fig. 2b) and a decrease in PSD length, notable in larger synapses (Con = 366±17 nm, DE = 335±12 nm; p<0.05, Mann-Whitney Test). The number of docked vesicles increased in smaller synapses and decreased in larger synapses (Con =3.14±0.11, DE = 3.91±0.11; p<0.01, Mann-Whitney Test). DE increased the area of the synaptic vesicle zone, notable in larger synapses, but not the overall presynaptic area (Synaptic vesicle zone, SVZ; Con = 0.17±0.9 μm^2^, DE = 0.25±0.3 μm^2^; p<0.01, Mann-Whitney Test, n = 264 and 219 synapses, respectively).

In contrast, brief LRx (2 hr) induced limited changes in synaptic ultrastructure. No change in synaptic cleft widths or PSD lengths were observed following LRx (cleft: DE = 19.0±0.2 nm, LRx = 19.2±0.1 nm; p=0.34, Mann-Whitney Test; PSD: DE = 335±12 nm, LRx = 328±7 nm; p=0.23; Mann-Whitney Test). However, the number of docked vesicles was decreased in smaller synapses and increased in larger synapses (DE = 3.91±0.11, LRx = 3.72±0.12; p<0.05, Mann-Whitney Test). Notably, LRx significantly reduced the area of the synaptic vesicle zone, but not overall presynaptic area (DE = 0.25±0.3 μm^2^, LRx = 0.19±0.1 μm^2^, p<0.01, Mann-Whitney Test, n = 219 and 320 synapses, respectively). Thus, DE induces widespread changes in synaptic ultrastructure, but changes induced by LRx are limited to parameters related to synaptic vesicle clustering and availability.

The declustering/clustering of synaptic vesicles by DE/LRx paralleled the declustering/clustering of presynaptic cytomatrix components observed following manipulations of synaptic activity in cultured hippocampal neurons (Glebov et al., 2016, 2017). To ask if DE/LRx impacted presynaptic molecular organization, we adapted a FRET-based intermolecular proximity assay to quantify *in situ* clustering of the cytomatrix protein bassoon (Bsn) in thalamic axons in layer 4 of adult mouse V1b (360±40 μm from dura). A validated monoclonal anti-Bsn primary antibody was followed by a pair of anti-mouse secondary antibodies conjugated to either a FRET acceptor or a FRET donor (1:1; Fig. 3a). Forster resonance energy transfer (FRET) distance is limited to ∼10 nm, therefore a change in FRET_Acceptor/Donor_ reports a change in intermolecular distance of the secondary antibodies.

**Fig. 3.**
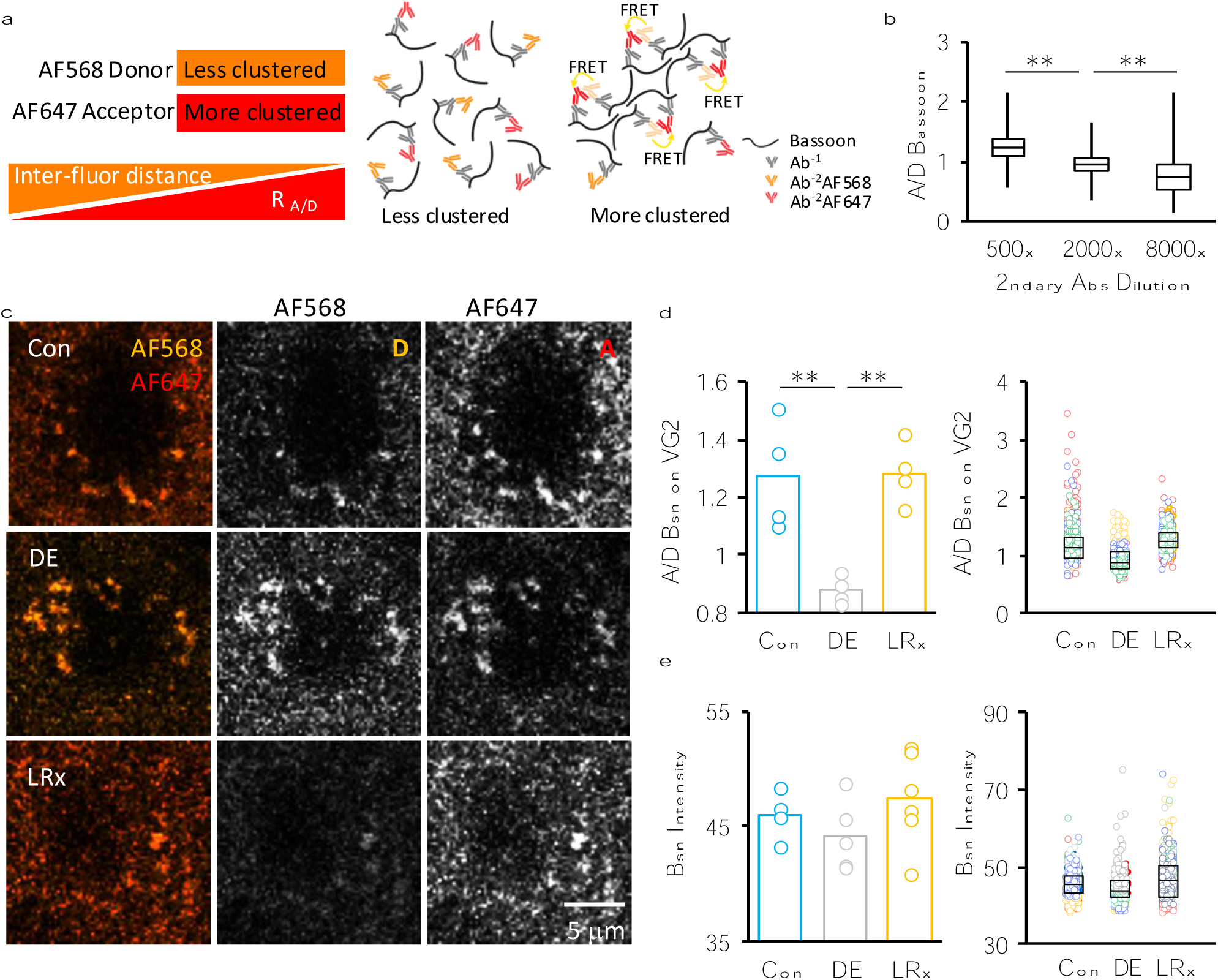
DE/LRx induce reversible changes in Bsn clustering in thalamic boutons of adult mouse V1b L4. a) Experimental logic of FRET-based proximity assay. b) FRET_A/D_ is decreased following serial dilution of secondary antibodies carrying FRET donor and FRET acceptor fluorophores. Box plots represent the median as a bar, 25th to 75th percentile as box, and max and min as whiskers (n = 2343, 2304, 970 puncta from 3 subjects each condition; One-way ANOVA, F = 1731, **p<0.01, *post hoc* Tukey). c) Representative examples of Bsn donor (orange, AF568) and acceptor (red, AF647) fluorescence in layer 4 of adult mouse V1b of Con, DE and LRx subjects. d) FRET_A/D_ values at VGluT2 puncta averaged by subject for Con, DE, LRx subjects (n = 4, 4, 4 subjects; One-way ANOVA, F = 13.0, **p<0.01, post hoc Tukey). Right: R_A/D_ of 100 puncta from each subject shown in randomly-assigned colors, overlayed with a box plot that represents the median as a bar, 25th to 75th percentile as box. e) Total Bsn immunofluorescence visualized with a single secondary antibody is unchanged by DE or LRx (One-way ANOVA, F = 1.0, p = 0.41. Right: Bsn immunofluorescence of 100 puncta from each subject shown in randomly-assigned colors, overlayed with a box plot that represents the median as a bar, 25th to 75th percentile as box. **p < 0.01 for *post hoc* tests.

The proximity assay is sensitive to intermolecular distance, as FRET_A/D_ decreased with serial dilution of the secondary antibody mix (500x, 1.24±0.004; 2000x, 0.95±0.003; 8000x, 0.78±0.01, n = 2343, 2304, 970 puncta, respectively. One-way ANOVA, F = 1731, p = 0.0001; *post hoc* Tukey, Fig. 3b). We then quantified the impact of DE/LRx on FRET_A/D_ in thalamic boutons identified by the presence of vesicular transporter 2 (VGluT2; (Nahmani and Erisir, 2005; Coleman et al., 2010)). DE significantly decreased FRET_A/D_, reporting an increase in intermolecular Bsn distance, and LRx significantly increased FRET_A/D_, reporting a decrease in intermolecular Bsn distance (Con =1.22±0.07, DE = 0.88±0.02, LRx = 1.28±0.03; n = 6, 4, 4 subjects, respectively, One-way ANOVA, F = 14.3, p < 0.001, *post hoc* Tukey; Fig. 3c, d). Neither DE nor LRx changed the total quantity of Bsn visualized with a single secondary antibody (Con = 45.9±0.9; DE = 44.2±1.4; LRx = 47.4±1.7; n = 6, 5, 6 subjects, respectively; One-way ANOVA, F = 1.0, p = 0.41; Fig. 3e). Thus, DE and LRx induce reversible changes in the molecular geometry of the presynaptic cytomatrix at thalamic boutons.

Clustering/unclustering of Bsn has been correlated with changes in the magnitude of evoked axonal calcium through P/Q type voltage gated calcium channels, a key determinant of neurotransmitter release (Glebov et al., 2017). We therefore asked if the magnitude of visually-evoked calcium transients at thalamic boutons in layer 4 was modified by DE/LRx (Fig. 4a). AAV5-Ef1a-DIO-Synaptophysin-GCaMP6s was delivered to dLGN in adult VGluT2-Cre mice 3 weeks prior to recording. Thalamic axons were imaged in V1b (average cortical depth: 320 μm). Calcium transients in axonal boutons were evoked by drifting, high-contrast, square wave gratings delivered to the dominant eye. Visually-responsive boutons were included for analysis if 1) peak ΔF/F value 0.25 - 2.0 sec after the onset of a visual stimulus in any direction exceeded 4 x the standard deviation (STD) of measurements at baseline (F_0_, −1.0 to 0 sec) for ≥ 40% trials and 2) the STD of ΔF/F during the rising phase of response (0.25 to 1.0 sec after visual stimulus onset) exceeded 2 x STD of baseline. dLGN boutons showed a preference for the orientation of the visual stimulus, as previously shown for thalamic boutons in layer 1 (Jaepel et al., 2017). ΔF/F of axonal bouton calcium transients evoked by the visual stimuli in the preferred orientation were compared in the same subject following DE, and LRx. DE decreased and LRx increased the magnitude of visually-evoked bouton calcium (Con = 1.065±0.03, DE = 0.48±0.03, LRx = 0.94±0.0; 1258, 2451, 1572 boutons from 4, 4, 4 subjects for Con, DE, and LRx, respectively; One-way ANOVA, F = 9.9, p = 0.005, *post hoc* Tukey; Fig. 4b-d). DE and LRx did not significantly impact the number of visually-responsive boutons (Con = 315±34, DE = 613±289, LRx = 393±89 boutons, n = 4 subjects, One-way ANOVA, F = 0.78, p = 0.49). Together, this demonstrates that DE/LRx in adulthood engages robust and reversible non-homeostatic changes in presynaptic function at feed-forward inputs from first order visual thalamus to visual cortex.

**Fig. 4.**
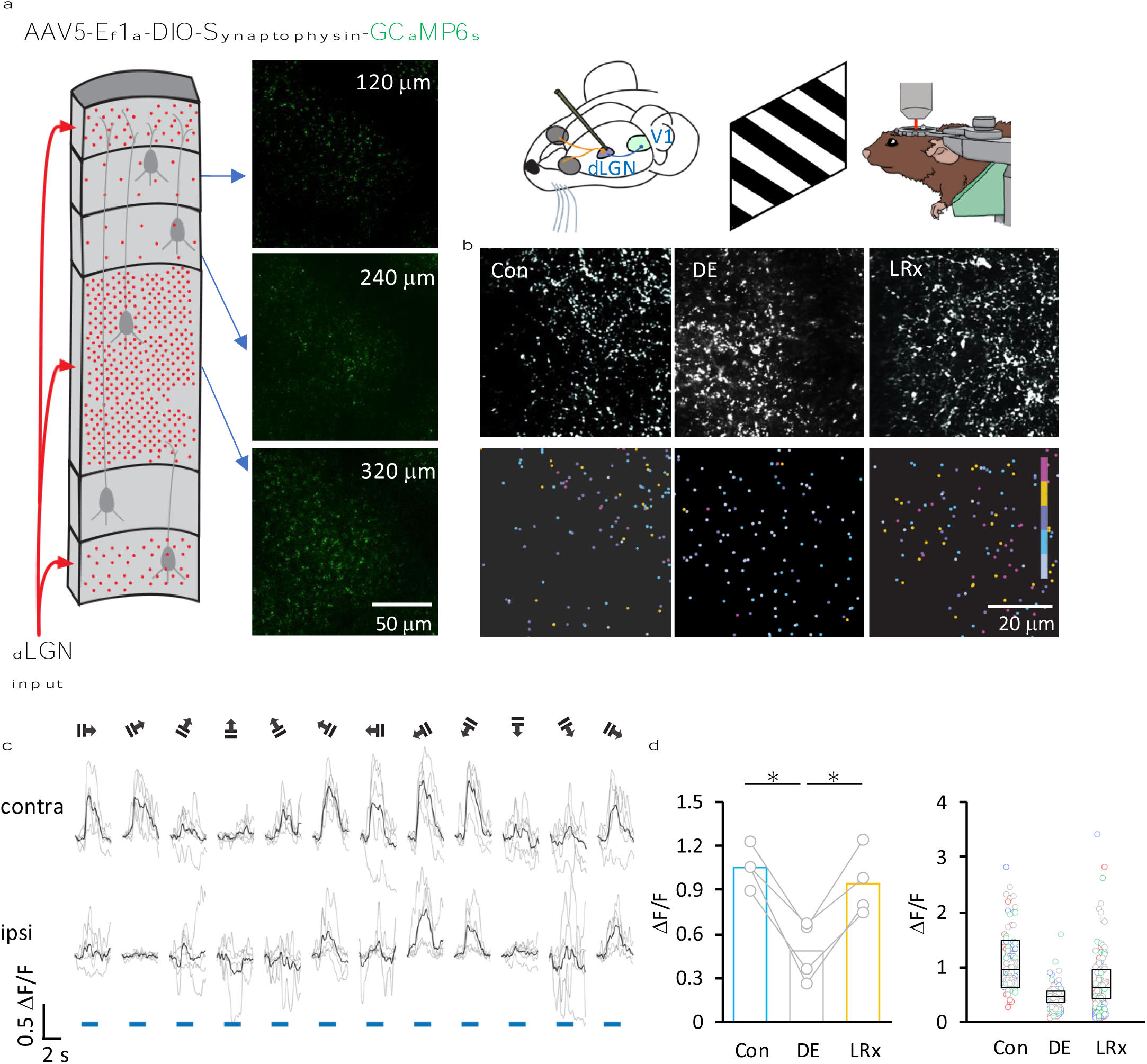
DE/LRx induce reversible changes in the magnitude of visually evoked calcium in dLGN boutons recorded in layer 4 of adult mouse V1b. a) AAV5-Ef1a-DIO-synaptophysin-GCaMP6s (750 μl) was delivered at 0.1 μl/min to dLGN: AP: 2.1 mm, ML: 2.2 mm, DV: 2.5 mm from bregma. 2P live imaging of axonal GCaMP in head-fixed awake adult mice. Cartoon depicting the abundance of thalamic axons throughout V1b. Single static images of axonal GCaMP6s at 3 depths acquired with low speed galvo scanning. b) Mean images after motion correction of 2P live axonal GCaMP6 imaging acquired with high speed resonant scanning (28Hz). Heat map depicting magnitude of visually-evoked ι1F/F for control (Con), DE and LRx subjects (gray—<0.3, blue—0.3 to 0.6, purple—0.6 to 0.9, orange—0.9 to 1.2, pink—>1.2). c) Representative examples of visually evoked GCaMP transients in single axonal boutons in response to 100% contrast, 0.05 cpd, square wave gratings of 12 directions drifting at 1 Hz. Blue bar = stimulus duration. d) Left: Visually-evoked GCaMP ι1F/F in thalamic boutons, evoked by stimulus of preferred orientation, and recorded in layer 4 of adult mouse V1b, averaged by subject (1258, 2451, 1572 boutons from n = 4, 4, 4 subjects for Con, DE, LRx. One-way ANOVA, F = 9.9, **p<0.01, post hoc Tukey, *p < 0.05 for *post hoc* tests). Right: ι1F/F for 30 boutons per subject shown in randomly-assigned colors, overlayed with a box plot that represents the median as a bar, 25th to 75th percentile as box.

## Discussion

Experience-dependent plasticity at synapses between first order thalamus and primary cortex was through to be constrained irreversibly by age. Indeed, the stability of TC synapses has been demonstrated many ways. Short term monocular deprivation does not change release probability of thalamic axons, and neither MD nor retinal activation decreases the overall firing output from dLGN neurons (Linden et al., 2009; Wang et al., 2013). However, we found that prolonged visual deprivation induced robust and reversible plasticity at thalamocortical synapses in adult mice that was non-homeostatic, and expressed as changes in presynaptic structure and function. Specifically, 1) Optogenetic control of dLGN boutons revealed that presynaptic release probability was reversibly decreased by DE and LRx. 2) Quantification of synaptic ultrastructure revealed that DE decreased and LRx increased the clustering and docking of synaptic vesicles in dLGN axonal boutons. 3) A FRET-based intermolecular proximity analysis revealed that the clustering of a presynaptic cytomatrix protein in thalamic axonal boutons was reversibly decreased by DE and LRx. 4) Intravital calcium imaging revealed that DE decreased and LRx increased the magnitude of visually evoked calcium in dLGN boutons in layer 4 of primary visual cortex. Thus, presynaptic depression of feed-forward drive from first order thalamus to primary visual cortex is engaged by prolonged activity deprivation and reversible upon recovery of afferent drive.

Our ultrastructural analysis of excitatory synapses in layer 4 demonstrated that fundamental parameters can be modified within a narrow range. A narrow range of synaptic cleft widths may be necessary to optimize trans-synaptic nanocolumns and limit neurotransmitter diffusion (Zuber et al., 2005; Savtchenko and Rusakov, 2007; Zheng et al., 2017; Cole and Reese, 2023). We adopted the area of the presynaptic terminal as a proxy for presynaptic origin, and ultrathin sections chosen randomly for quantification to avoid sampling bias (Schoonover et al., 2014; Bopp et al., 2017). The interpretation that the largest 15% of synaptic profiles are enriched for TC synapses mirrors the estimated abundance of thalamic boutons in layer 4 (Nahmani and Erisir, 2005; Coleman et al., 2010). Importantly, DE/LRx induces opposing changes in ultrastructure, consistent with non-homeostatic changes at larger, thalamocortical synapses and homeostatic changes at smaller, corticocortical synapses.

Thalamic axonal boutons do not express the synaptic vesicle-binding protein synapsin, and are therefore predicted to employ different strategies to regulate synaptic vesicle organization (Owe et al., 2013). We found that in adult thalamocortical synapses in situ, Bsn declustering correlates with reduced visually-evoked calcium in axonal boutons and decreased neurotransmitter release probability. In contrast, at corticocortical synapses, bassoon deletion reduces vesicle release competency and dysregulates PKA-dependent phosphorylation (Milovanovic et al., 2018; Montenegro-Venegas et al., 2022). Action potential suppression in cultured hippocampal neurons declusters Bsn, increases the density of active zone Cav2.1 channels and increases action potential-evoked vesicle release (Glebov et al., 2017).

Homeostatic changes to compensate for the reduction in synaptic drive is a prevailing response to prolonged activity deprivation. In some cases, prolonged activity suppression can also induce heterogeneous changes in pre- and postsynaptic function, resulting in fewer but stronger excitatory synapses (Mitra et al., 2012; Wise et al., 2024). However, we observed no difference in the number of visually-responsive axonal boutons and consistent modification in neurotransmitter release probability following DE/LRx. In contrast to limitations of plasticity from first order thalamus to layer 4, plasticity in synapses between the dLGN shell and layer 1 have been reported in adult (Sohn et al., 2022) including MD-induced changes in the amplitude of visually-evoked calcium in dLGN axonal boutons synapsing in layer 1 of primary visual cortex (Jaepel et al., 2017). Monocular deprivation in adults induces an increase in the visually evoked spiking in response to the non-deprived eye in both dLGN and V1 neurons. Notably pharmacological inhibition of primary visual cortex blocked this response to monocular deprivation in juveniles but not adults, suggesting feedback from cortex to thalamus is also constrained by age (Qin et al., 2023).

To our knowledge, this is the first demonstration of plasticity at synapses between first order thalamus and primary sensory cortex in adults. Unexpectedly, these synapses express a non-homeostatic response with a presynaptic locus of expression in response to prolonged visual deprivation. A recent report heralding the potential for activity-dependent regulation of these synapses demonstrated transcranial low-intensity focused ultrasound can induce a long-term depression of thalamic inputs to layer 4 A recent report signaling the potential for activity-dependent regulation of these synapses by demonstrating transcranial low-intensity focused ultrasound can induce a long-term depression of thalamic inputs to layer 4(Mesik et al., 2024). Our results support the idea that uncommon changes in synaptic activity may be required to engage plasticity at adult TC synapses. The presynaptic plasticity mechanism with a high threshold for engagement described here would maintain synapse stability across a range of activity, and allow adaptation only in extraordinary conditions.

## Materials and methods

### Subjects

C57BL/6J mice were purchased from Jackson Laboratory (Bar Harbor, ME, USA). Equal numbers of adult (>P90) males and females were used. Mice were raised in a 12:12 h light/dark cycle. All procedures conformed to the guidelines of the University of Maryland and Johns Hopkins University Institutional Animal Care and Use Committees. Control experiments were performed (or subjects were sacrificed) 6 h into the light phase.

### V1 slice preparation and optogenetic thalamocortical synapse isolation

AAV5-CaMKIIa-hChR_2_(H134R)-mCherry (addgene, Cat# 26975-AAV5, 1.4 x 10^13^ GS/mL, 300 μl) was delivered to adult C57BL/6J mouse dLGN (from Bregma AP: 2.3 mm, ML: 2.0 mm, DV: 2.4 mm) 3 weeks prior to recording at >postnatal day 90 (>P90). Mice were anesthetized using isoflurane vapors; after disappearance of the corneal reflex mice were transcardially perfused with ice-cold dissection buffer containing (in mM): 212.7 sucrose, 5 KCl, 1.25 NaH_2_PO_4_, 10 MgCl_2_, 0.5 CaCl_2_, 26 NaHCO_3_, and 10 dextrose, saturated with 95% O_2_/5% CO_2_ (pH 7.4). The brain was rapidly removed, immersed in ice-cold dissection buffer and sectioned (300 µm) using a vibratome (Leica VT1200S). Visual cortical slices were transferred to artificial cerebrospinal fluid (ACSF), incubated at 30 °C for 30 min, and room temperature for 30 min then transferred to the recording chamber. ACSF was similar to dissection buffer except that sucrose was replaced by 124 mM NaCl, MgCl_2_ was lowered to 1 mM, and CaCl_2_ was raised to 2 mM. Visualized whole-cell recordings were made from pyramidal neurons in layer 4 of V1 with glass pipettes (3-5 MΩ) filled with (in mM) 8 KCl, 125 cesium gluconate, 10 HEPES, 1 EGTA, 4 MgATP, 0.5 NaGTP, and 5 QX-314 (pH 7.2 to 7.3 and 280 to 295 mOsm). Biocytin (1 mg/ml) was added to the internal solution for *post hoc* cell identification. ChR_2_-containing axon terminals were activated using a 470 nm wavelength LED (3 ms duration; Thorlabs) through a 40x objective lens. EPSCs were recorded in voltage–clamp at −70 mV.

Sr^2+^-desynchronized miniature EPSCs (Sr^2+^ mEPSCs, Fig. 1c, d) were evoked at the light intensity evoking a response in 50% of trials (470 nm, 3 msec) following replacement of Ca^2+^ with 4 mM Sr^2+^ and raising Mg^2+^ concentration to 4 mM (Dodge Jr. et al., 1969). To ensure that light-evoked responses are monosynaptic, recordings were performed in the presence of 1 μM TTX and 100 μM 4-AP (Petreanu et al., 2009; Whitt et al., 2022). Only cells with series resistance <25 MΩ and <25% variation over the experiment were included. Data was filtered at 4 kHz and digitized at 10 kHz using Igor Pro (Wave Metrics).

To quantify the response to repetitive optogenetic stimulation (Fig. 1e-j), 15 light pulses were delivered at 25 Hz. The minimal light intensity that evoked a response in 100% of trials. EPSCs were recorded in the presence of 100 μM APV and 20 μM bicuculine. The shape of the EPSC was examined, and cells were excluded from analysis if the rising phase exhibited any inflection points, which could indicate a polysynaptic response. The paired pulse ratio was calculated as 2^nd^ EPSC/1^st^ EPSC. The normalized EPSC at steady state was calculated as the mean of 11^th^ to 15^th^ EPSC. The release probability and the recovery rate were calculated as described in (Bridi et al., 2020) based on (Wesseling and Lo, 2002). Briefly, equations (2) and (3) were derived from equation (1) as shown below:

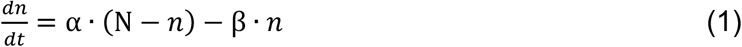

Where α denotes the recovery rate, β denotes the release probability, *n* represents the vesicles available for release, and *N* represents the total number of vesicles in the read releasable pool.

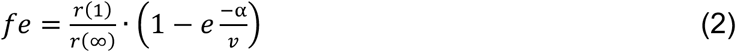

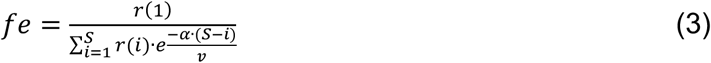

Where *fe* is the initial release probability, *r*(1) is the amount of neurotransmitter released by the first stimulus in the train, *r*(∞) represents the release at steady-state during the train, *v* is the stimulation frequency, *S* is the total number of stimuli in the train, *i* denotes the *i*-th stimulus, *r*(*i*) is the release corresponding to the *i*-th stimulus, and *r*(*i*)/*r*(1) represents the normalized amplitude of the *i*-th EPSC in the train. The values for *fe* and *α* were determined by solving equations (2) and (3).

### Biocytin processing

After recordings, slices were fixed in 4% paraformaldehyde overnight at 4°C. Slices were rinsed 2 × 10 min in 0.1 M phosphate buffer (PB) containing 19 mM NaH_2_PO_4_ · H_2_O and 81 mM Na_2_HPO_4_ at room temperature and permeabilized in 2% Triton X-100 in 0.1 mM PB for 1 h. Slices were then incubated in avidin-Alexa Fluor 633 conjugate diluted 1:2000 in 1% Triton X-100/0.1 M PB overnight at 4°C in the dark. After the incubation, slices were washed 2 × 10 min in 0.1 M PB, mounted on glass slides, and allowed to dry overnight in the dark. Slides were cover-slipped with Prolong Anti-fade (Invitrogen) mounting medium and sealed with nail polish. Images were taken using a Zeiss LSM 700 confocal microscope.

### Electron Microscopy

Mice were transcardially perfused with 2.5% glutaraldehyde and 4% paraformaldehyde (PFA) in phosphate buffered saline (PBS, pH 7.4). Brains were post-fixed overnight and stored in 0.01% NaN_3_ in PBS at 4°C. Coronal sections (100 microns) were made using a Leica vibratome (VT1000), and V1b and the hippocampal commissure were isolated. Sections were rinsed in 0.1M sodium cacodylate buffer. The following processing steps were then carried out using the variable wattage Pelco BioWave Pro microwave oven (Ted Pella, Inc., Redding, CA): post-fixation in 1% osmium tetroxide made in 0.1 M sodium cacodylate buffer, rinse in double distilled water (DDW), 2% (aq.) uranyl acetate enhancement, DDW rinse, ethanol dehydration series up to 100% ethanol and propylene oxide, followed by an Embed-812 resin (Electron Microscopy Sciences, Hatfield, PA) infiltration series up to 100% resin. The epoxy resin was polymerized for 20 h in an oven at 60°C. Ultra-thin sections were cut on a Leica EM-UC7 Ultramicrotome (90 nm). Thin sections were placed on 200 mesh copper grids and post-stained with uranyl acetate and lead citrate. Images were collected 360±40 μm from dura. Imaging was accomplished using a JEOL-1400 Transmission Electron Microscope operating at 80kV and an AMT BioSprint-29 camera at 25000 times magnification at 0.37 nm/pixel resolution. All synaptic profiles with high contrast boundaries delineating the presynaptic membrane, active zone, post-synaptic membrane and post synaptic density were quantified. Image acquisition and analysis were performed blind.

### FRET-based molecular proximity assay

Subjects were anesthetized with 4% isoflurane in O_2_ and perfused with PBS followed by 4% PFA in PBS. The brains were post-fixed in 4% PFA for 24 h, followed by 30% sucrose for 24 h, and cryo-protectant solution for 24 h (0.58 M sucrose, 30% (v/v) ethylene glycol, 3 mM sodium azide, 0.64 M sodium phosphate, pH 7.4). Coronal sections (40 μm) were cut on a Leica vibratome (VT1000). Sections were blocked with 4% normal goat serum (NGS) containing 0.4% TritonX-100 and 0.1% Tween-20 in PBS for 30 min. The primary antibody was presented in blocking solution overnight at 4°C, followed by secondary antibodies for 6 hours.

The following antibodies/dilutions were used: Monoclonal mouse anti-Bsn (NeuroMab) RRID: AB_2895685, 1:1000, goat anti-mouse IgG Alexa-568 conjugated (Thermo Fisher Scientific) RRDI: AB_2534072, 1:1000, goat anti-mouse IgG Alexa-647 conjugated (Thermo Fisher Scientific) RRID: AB_2535804, 1:1000, unless the dilution is specified. FRET experiments were performed with the anti-Bsn monoclonal mouse antibody and a 50:50 mixture of the two secondary antibodies.

Images were acquired on a Leica SP5x confocal microscope with a 60X Oil lens (HCX PL APO CS, NA = 1.4) at Ex = 543 nm, with bandpass filters for Alexa-568: 560 – 615 nm and Alexa-647: 650 – 720 nm. Bsn puncta were analyzed in single Z-section images in Fiji (NIH) by applying the threshold function (auto threshold + 20) and size exclusion (0.2–2.0 μm^2^) using the *analyze particles* function. Image acquisition and analysis were performed blind.

### Intravital imaging of axonal GCaMP

Axonal GCaMP6s was delivered to dLGN (from Bregma AP: 2.1 mm, ML: 2.2 mm, DV: 2.5 mm) via AAV5-Ef1a-DIO-Synaptophysin-GCaMP6s (addgene, Cat# 105715-AAV5, 2.0 x 10^13^ GS/ml) injected via a Hamilton syringe attached to a Microsyringe Pump Controller (World Precision Instruments) at a rate of 100 nl/min (total volume of 750 μl) to adult (>postnatal day 90, P>90) VGluT2-Cre mice (Jax strain 028863).

A cranial window consisting of two 3 mm diameter coverslips glued with optical adhesive (Norland71, Edmund Optics) to a 5 mm diameter coverslip was implanted. The gap between the skull and glass was sealed with silicone elastomer (Kwik-Sil). Instant adhesive Loctite 454 (Henkel) was used to adhere an aluminum head post to the skull, and to cover the exposed skull. Black dental cement (iron oxide powder, AlphaChemical mixed with white powder, Dentsply) was used to coat the surface to minimize light reflections. Subjects were imaged ≥ 3 weeks of recovery.

Awake subjects were placed in a holding tube and immobilized by a head post clamp. Prior to the first imaging session, the subjects were habituated to the holding tube at least twice for >30 min. A shield was placed around the gap between the cranial window and the objective lens to block light contamination from the visual stimuli during imaging. An Olympus FVMPE-RS Multiphoton Laser Scanning Microscope controlled by Fluoview software with a 25x NA 1.05 water immersion objective lens was used to acquire time lapse fluorescence images. An Insight X3-OL laser was tuned to 940 nm for imaging GCaMP (Ex max 490 nm). Fluorescence emission was detected through a dichroic mirror (495-540 nm). The field of view was 169.71 μm x 169.71 μm (512×512 pixels), 360±40 μm from the brain surface. Minimum laser power (59 mW) and PMT gain (400) necessary to image at this depth was used. We observed no photobleaching with these imaging parameters. Images of static representations are average of 2 acquired in galvanometer scanning mode, images of visually evoked GCaMP ΔF/F were acquired in resonant scanning mode at 28 Hz. Using the Suite2P package (Pachitariu et al., 2017), movement artifacts was corrected using the average intensity of the full image stack as a template. Region of interests (ROIs) were detected by axon/bouton mode (correction factor, 0.7).

To evoke visual responses in boutons, subjects received monocular visual stimuli controlled by PsychToolBox plugin in MATLAB (random order of 5 repeats of 12 directions of square wave gratings drifting at 1 Hz at 0.05 cycle/degree (cpd), 100% contrast for 2.5 s interleaved with 2.5 s intensity-matched (28 cd/m^2^) grey scale, delivered by a 23” display (Acer LCD Monitor) 28 cm in front of the eyes). Calculation of ΔF/F = (F-F0)/F0, where F corresponds to the fluorescence intensity at a given time point and F0 corresponds to mean fluorescent intensity during 1 s before the visual stimulus onset. A bouton was defined as visually responsive if the peak ΔF/F value 0.25 to 2.0 sec after visual stimulation onset at any direction exceeded 4 x STD of baseline (F0, −1.0 to 0 sec) for ≥ 40% of trials during either contra or ipsi eye stimulation, and if the STD of ΔF/F during the rising phase of response (0.25 to 1.0 sec after visual stimulus onset) exceeds 2 x STD of the baseline. All layer 4 TC boutons that were visually-responsive were included in our analysis. We employed a population analysis to control for difference in bouton identity across recording conditions.

### Statistics

An unpaired two-tailed Student’s T-test was used to determine the significance between two independent experimental groups. A Mann-Whitney test was applied to EM analyses due to the abnormal distribution caused by the bimodal distribution of presynaptic area. A one-way ANOVA was performed to determine the significance between three or more independent experimental groups. Kruskal-Wallis was used if an abnormal distribution was predicted. For spike train experiments, a two-way ANOVA was performed.

## Acknowledgement

This work was supported by R01EY016431 (EMQ) EY025922 (AK and EMQ), R01EY12124 (AK), and NICHD IRP (LD). We thank Dr. Ji Liu for technical assistance on 2P imaging acquisition and analyses.

## References

Antonini A, Stryker MP (1993) Rapid Remodeling of Axonal Arbors in the Visual Cortex. Science (1979) 260:1819–1821.

Antonini A, Stryker MP (1996) Plasticity of geniculocortical afferents following brief or prolonged monocular occlusion in the cat. Journal of Comparative Neurology 369:64–82.

Bopp R, Holler-Rickauer S, Martin KAC, Schuhknecht GFP (2017) An ultrastructural study of the thalamic input to layer 4 of primary motor and primary somatosensory cortex in the mouse. Journal of Neuroscience 37:2435–2448.

Bridi MCD, de Pasquale R, Lantz CL, Gu Y, Borrell A, Choi S-Y, He K, Tran T, Hong SZ, Dykman A, Lee H-K, Quinlan EM, Kirkwood A (2018) Two distinct mechanisms for experience-dependent homeostasis. Nat Neurosci 21:843–850.

Bridi MS, Shin S, Huang S, Kirkwood A (2020) Dynamic Recovery from Depression Enables Rate Encoding in Inhibitory Synapses. iScience 23:100940.

Chokshi V, Gao M, Grier BD, Owens A, Wang H, Worley PF, Lee HK (2019) Input-Specific Metaplasticity in the Visual Cortex Requires Homer1a-Mediated mGluR5 Signaling. Neuron 104:736–748.e6.

Chung S, Jeong JH, Ko S, Yu X, Kim YH, Isaac JTR, Koretsky AP (2017) Peripheral Sensory Deprivation Restores Critical-Period-like Plasticity to Adult Somatosensory Thalamocortical Inputs. Cell Rep 19:2707–2717.

Cole AA, Reese TS (2023) Transsynaptic Assemblies Link Domains of Presynaptic and Postsynaptic Intracellular Structures across the Synaptic Cleft. Journal of Neuroscience 43:5883–5892.

Coleman JE, Nahmani M, Gavornik JP, Haslinger R, Heynen AJ, Erisir A, Bear MF (2010) Rapid Structural Remodeling of Thalamocortical Synapses Parallels Experience-Dependent Functional Plasticity in Mouse Primary Visual Cortex. J Neurosci 30:9670–9682.

Cooper LN, Bear MF (2012) The BCM theory of synapse modification at 30: interaction of theory with experiment. Nat Rev Neurosci 13:798–810.

Crair MC, Malenka RC (1995) A critical period for long-term potentiation at thalamocortical synapses. Nature 375:325–328.

Dodge Jr. FA, Miledi R, Rahamimoff R (1969) Strontium and quantal release of transmitter at the neuromuscular junction. J Physiol 200:267–283.

Duffy KR, Lingley AJ, Holman KD, Mitchell DE (2016) Susceptibility to monocular deprivation following immersion in darkness either late into or beyond the critical period. Journal of Comparative Neurology 524:2643–2653.

Eaton NC, Sheehan HM, Quinlan EM (2016) Optimization of visual training for full recovery from severe amblyopia in adults. Learning and Memory 23:99–103.

Feldman DE, Nicoll RA, Malenka RC, Isaac JTR (1998) Long-term depression at thalamocortical synapses in developing rat somatosensory cortex. Neuron 21:347–357.

Glebov OO, Cox S, Humphreys L, Burrone J (2016) Neuronal activity controls transsynaptic geometry. Sci Rep 6:1–11.

Glebov OO, Jackson RE, Winterflood CM, Owen DM, Barker EA, Doherty P, Ewers H, Burrone J (2017) Nanoscale Structural Plasticity of the Active Zone Matrix Modulates Presynaptic Function. Cell Rep 18:2715–2728.

Goel A, Jiang B, Xu LW, Song L, Kirkwood A, Lee H-K (2006) Cross-modal regulation of synaptic AMPA receptors in primary sensory cortices by visual experience. Nat Neurosci 9:1001–1003.

Goel A, Lee H-K (2007) Persistence of Experience-Induced Homeostatic Synaptic Plasticity through Adulthood in Superficial Layers of Mouse Visual Cortex. Journal of Neuroscience 27:6692–6700.

He HY, Hodos W, Quinlan EM (2006) Visual deprivation reactivates rapid ocular dominance plasticity in adult visual cortex. Journal of Neuroscience 26:2951–2955.

He H-Y, Ray B, Dennis K, Quinlan EM (2007) Experience-dependent recovery of vision following chronic deprivation amblyopia. Nat Neurosci 10:1134–1136.

Isaac JTR, Crair MC, Nicoll RA, Malenka RC (1997) Silent synapses during development of thalamocortical inputs. Neuron 18:269–280.

Jaepel J, Hübener M, Bonhoeffer T, Rose T (2017) Lateral geniculate neurons projecting to primary visual cortex show ocular dominance plasticity in adult mice. Nat Neurosci 20:1708–1714.

Jeon BB, Fuchs T, Chase SM, Kuhlman SJ (2022) Visual experience has opposing influences on the quality of stimulus representation in adult primary visual cortex. Elife 11:1–18.

Jiang B, Trevino M, Kirkwood A (2007) Sequential Development of Long-Term Potentiation and Depression in Different Layers of the Mouse Visual Cortex. Journal of Neuroscience 27:9648–9652.

Jie H, Petrus E, Pothayee N, Koretsky AP (2025) Reactivated thalamocortical plasticity alters neural activity in sensory-motor cortex during post-critical period. Prog Neurobiol 247.

Lee HK, Kirkwood A (2019) Mechanisms of Homeostatic Synaptic Plasticity in vivo. Front Cell Neurosci 13:1–7.

Linden ML, Heynen AJ, Haslinger RH, Bear MF (2009) Thalamic activity that drives visual cortical plasticity. Nat Neurosci 12:390–392.

Mainardi M, Landi S, Gianfranceschi L, Baldini S, De Pasquale R, Berardi N, Maffei L, Caleo M (2010) Environmental enrichment potentiates thalamocortical transmission and plasticity in the adult rat visual cortex. J Neurosci Res 88:3048–3059.

Mesik L, Parkins S, Severin D, Grier BD, Ewall G, Kotha S, Wesselborg C, Moreno C, Jaoui Y, Felder A, Huang B, Johnson MB, Harrigan TP, Knight AE, Lani SW, Lemaire T, Kirkwood A, Hwang GM, Lee HK (2024) Transcranial Low-Intensity Focused Ultrasound Stimulation of the Visual Thalamus Produces Long-Term Depression of Thalamocortical Synapses in the Adult Visual Cortex. Journal of Neuroscience 44:1–12.

Milovanovic D, Wu Y, Bian X, De Camilli P (2018) A liquid phase of synapsin and lipid vesicles. Science (1979) 361:604–607.

Min SH, Wang Z, Chen MT, Hu R, Gong L, He Z, Wang X, Hess RF, Zhou J (2023) Metaplasticity: Dark exposure boosts local excitability and visual plasticity in adult human cortex. Journal of Physiology 601:4105–4120.

Mitra A, Mitra SS, Tsien RW (2012) Heterogeneous reallocation of presynaptic efficacy in recurrent excitatory circuits adapting to inactivity. Nat Neurosci 15:250–257.

Montenegro-Venegas C, Guhathakurta D, Pina-Fernandez E, Andres-Alonso M, Plattner F, Gundelfinger ED, Fejtova A (2022) Bassoon controls synaptic vesicle release via regulation of presynaptic phosphorylation and cAMP. EMBO Rep 23:1–20.

Montey KL, Quinlan EM (2011) Recovery from chronic monocular deprivation following reactivation of thalamocortical plasticity by dark exposure. Nat Commun 2.

Murase S, Lantz CL, Quinlan EM (2017) Light reintroduction after dark exposure reactivates plasticity in adults via perisynaptic activation of MMP-9. Elife 6.

Nahmani M, Erisir A (2005) VGluT2 immunochemistry identifies thalamocortical terminals in layer 4 of adult and developing visual cortex. J Comp Neurol 484:458–473.

Owe SG, Erisir A, Heggelund P (2013) Terminals of the major thalamic input to visual cortex are devoid of synapsin proteins. Neuroscience 243:115–125.

Pachitariu M, Stringer C, Dipoppa M, Schröder S, Rossi LF, Dalgleish H, Carandini M, Harris KD (2017) Suite2p: beyond 10,000 neurons with standard two-photon microscopy. bioRxiv. Bioarxiv 20:2017.

Petreanu L, Mao T, Sternson SM, Svoboda K (2009) The subcellular organization of neocortical excitatory connections. Nature 457:1142–1145.

Petrus E, Isaiah A, Jones AP, Li D, Wang H, Lee HK, Kanold PO (2014) Crossmodal Induction of Thalamocortical Potentiation Leads to Enhanced Information Processing in the Auditory Cortex. Neuron 81:664–673.

Petrus E, Rodriguez G, Patterson R, Connor B, Kanold PO, Lee H-K (2015) Vision loss shifts the balance of feedforward and intracortical circuits in opposite directions in mouse primary auditory and visual cortices. Journal of Neuroscience 35:8790–8801.

Qin Y, Ahmadlou M, Suhai S, Neering P, de Kraker L, Heimel JA, Levelt CN (2023) Thalamic regulation of ocular dominance plasticity in adult visual cortex. Elife 12:1–17.

Sale A, Maya Vetencourt JF, Medini P, Cenni MC, Baroncelli L, De Pasquale R, Maffei L (2007) Environmental enrichment in adulthood promotes amblyopia recovery through a reduction of intracortical inhibition. Nat Neurosci 10:679–681.

Savtchenko LP, Rusakov DA (2007) The optimal height of the synaptic cleft. Proc Natl Acad Sci U S A 104:1823–1828.

Schoonover CE, Tapia JC, Schilling VC, Wimmer V, Blazeski R, Zhang W, Mason CA, Bruno RM (2014) Comparative strength and dendritic organization of thalamocortical and corticocortical synapses onto excitatory layer 4 neurons. Journal of Neuroscience 34:6746–6758.

Sohn J, Suzuki M, Youssef M, Hatada S, Larkum ME, Kawaguchi Y, Kubota Y (2022) Presynaptic supervision of cortical spine dynamics in motor learning. Available at: https://www.science.org.

Stodieck SK, Greifzu F, Goetze B, Schmidt KF, Löwel S (2014) Brief dark exposure restored ocular dominance plasticity in aging mice and after a cortical stroke. Exp Gerontol 60:1–11.

Wang L, Kloc M, Gu Y, Ge S, Maffei A (2013) Layer-specific experience-dependent rewiring of thalamocortical circuits. Journal of Neuroscience 33:4181–4191.

Wen W, Turrigiano GG (2024) Keeping Your Brain in Balance: Homeostatic Regulation of Network Function. Annu Rev Neurosci 47:41–61.

Wesseling JF, Lo DC (2002) Limit on the role of activity in controlling the release-ready supply of synaptic vesicles. Journal of Neuroscience 22:9708–9720.

Whitt JL, Ewall G, Chakraborty D, Adegbesan A, Lee R, Kanold PO, Lee HK (2022) Visual Deprivation Selectively Reduces Thalamic Reticular Nucleus-Mediated Inhibition of the Auditory Thalamus in Adults. Journal of Neuroscience 42:7921–7930.

Wiesel TN, Hubel DH (1963) Responses in Striate Deprived of Vision Cortex of One Eye. J Neurophysiol 26:1003–1017.

Wise DL, Escobedo-Lozoya Y, Valakh V, Gao EY, Bhonsle A, Lei QL, Cheng X, Greene SB, Van Hooser SD, Nelson SB (2024) Prolonged Activity Deprivation Causes Pre-and Postsynaptic Compensatory Plasticity at Neocortical Excitatory Synapses. eNeuro 11:1–12.

Yu X, Chung S, Chen DY, Wang S, Dodd SJ, Walters JR, Isaac JTR, Koretsky AP (2012) Thalamocortical Inputs Show Post-Critical-Period Plasticity. Neuron 74:731–742.

Zheng K, Jensen TP, Savtchenko LP, Levitt JA, Suhling K, Rusakov DA (2017) Nanoscale diffusion in the synaptic cleft and beyond measured with time-resolved fluorescence anisotropy imaging. Sci Rep 7:1–10.

Zuber B, Nikonenko I, Klauser P, Muller D, Dubochet J (2005) The mammalian central nervous synaptic cleft contains a high density of periodically organized complexes. Proc Natl Acad Sci U S A 102:19192–19197.

